# A pathological phosphorylation pattern enhances tau cooperativity on microtubules and facilitates tau filament assembly

**DOI:** 10.1101/2025.01.29.635117

**Authors:** Xiangyu Fan, Kyoko Okada, Henry Lin, Kassandra M. Ori-McKenney, Richard J. McKenney

## Abstract

Phosphorylation plays a crucial role in both normal and disease processes involving the microtubule-associated protein tau. Physiologically, phosphorylation regulates tau’s subcellular localization within neurons and is involved in fetal development and animal hibernation. However, abnormal phosphorylation of tau is linked to the formation of neurofibrillary tangles (NFTs) in various human tauopathies. Interestingly, the patterns of tau phosphorylation are similar in both normal and abnormal processes, leaving unclear whether phosphorylated tau retains its functional role in normal processes. The relationship between tau phosphorylation and NFT assembly in tauopathies is also still debated. To address these questions, we investigated the effects of tau phosphorylation on microtubule binding, cooperative protein envelope formation, and NFT filament assembly relevant to tauopathies. Consistent with previous results, our findings show that tau phosphorylation decreases tau’s overall affinity for microtubules, but we reveal that phosphorylation more dramatically impacts the cooperativity between tau molecules during tau envelope formation along microtubules. Additionally, we observed that the specific pattern of phosphorylation, rather than overall phosphorylation level, strongly impacts the assembly of tau filaments *in vitro*. Our results reveal new insights into how tau phosphorylation impacts tau’s physiological roles on microtubules and its pathoconversion into NFTs.

## Introduction

Tau was first identified as a microtubule-associated protein (MAP) in 1975^1^, and was initially thought to play a key role in regulating microtubule dynamics by promoting polymerization and stability^2,3^. Further research revealed that tau also participates in many cellular processes, including the activation of MAPK (mitogen-activated protein kinase)^4^, the regulation of synaptic function^5–7^, the protection of RNA and DNA in nucleus^8–11^, and the regulation of protein synthesis^12^. More recently, we and others have shown that tau plays a crucial role in regulating microtubule-based activities by forming cooperative protein “envelopes” along microtubules, which act as selectively permeable barriers for other MAPs^13,14^, impacting microtubule-based cargo transport^15,16^. In this process, these cohesive tau envelopes work as microtubule remodeler to compact the lattice of the microtubule^17,18^. The research interest in tau rose sharply after 1985 when tau was identified as a primary constituent of the neurofibrillary tangles (NFTs) in Alzheimer’s disease (AD) patient’s brain^19–22^, as well as other neurodegenerative diseases, such as chronic traumatic encephalopathy (CTE)^23,24^, progressive supranuclear palsy (PSP) and Pick’s disease (PiD)^25^. NFTs are composed of paired-helical filaments (PHFs) and straight filaments (SHs), which are polymers of tau protein that has undergone a structural transition from an intrinsically disordered state to a highly ordered cross-β structure. Recent high-resolution structural studies have elucidated the atomic arrangement of these tau filaments, providing novel insights into this structural transformation^26–29^. However, the molecular mechanism underlying the conversion of tau protein into this highly ordered state remains poorly understood and is the subject of ongoing research. The accumulation of NFTs in tauopathies is closely linked to neuronal cell death and resulting cognitive decline, ultimately leading to dementia and death. The link between tau’s physiological roles and conversion to pathological states remains undefined and stands as one of the great mysteries obfuscating a fuller understanding of the dementia process.

Phosphorylation of tau plays a key role in mediating tau’s physiological and pathological roles. Tau phosphorylation is a hallmark of neurofibrillary tangles (NFTs) in various tauopathies^20–25^. Research has consistently shown that multi-site phosphorylation of tau alters its conformation, reduces its affinity for microtubules, and promotes aggregation^30–39^. Additionally, abnormal tau phosphorylation leads to mislocalization of tau from the axon to the cytosol within neurons^40^. Interestingly, phosphorylation also plays a crucial role in normal physiological processes, such as animal hibernation^41–43^, and fetal development^44–46^. In hibernating animals, tau protein exhibits elevated phosphorylation levels during torpor, a state characterized by reduced metabolic rate and body temperature. This phenomenon is likely influenced by the altered activity of kinases and phosphatases, which are sensitive to the low metabolic and thermal conditions. Following arousal from hibernation, the phosphorylation levels of tau protein return to normal. Notably, this hibernation-arousal cycle does not result in brain damage, suggesting that the brain is resilient to the changes in tau protein phosphorylation that occur during torpor^41–43,47^. In fetal brain development, tau is highly phosphorylated at multiple sites, but this phosphorylation level returns to a steady state after maturation due to increased phosphatase activity in adult brains^44^. Interestingly, the fetal tau shows similar phosphorylation level with the tau from AD brain, but with no signs of aggregation or NFT formation^39,45,46,48^. Furthermore, physiological tau phosphorylation is typically thought to be a reversible process, unlike pathological tau phosphorylation^41–43^.

The physiological functions of phosphorylated tau, including its regulation of microtubule-based activities, remain poorly understood. Furthermore, the impact of phosphorylation on the interactions between tau molecules on microtubules and during the assembly of pathological NFTs is still unclear. To address these knowledge gaps, we employed tau proteins with varying patterns of phosphorylation and investigated the effects of phosphorylation on microtubule binding, cooperative tau envelope formation, oligomerization, and filament assembly relevant to tauopathies. Our findings reveal that tau phosphorylation alters the single molecule kinetics and cooperative interactions that underly tau envelope formation along microtubules, but that phosphorylated tau can still bind and compact the microtubule lattice, albeit with reduced affinity. We discovered that human tau protein produced in insect cells, which exhibits a phosphorylation pattern similar to that found in Alzheimer’s disease brains^49^, alters cooperative interactions between tau molecules and impacts cooperative tau envelope assembly. Additionally, our results show that phosphorylation promotes oligomerization of full-length tau molecules in solution. Finally, *in vitro* co-assembly assays revealed that pathological phosphorylation patterns greatly facilitate the assembly of tau proteins into NFT-like filaments under conditions that promote filament structures found in chronic traumatic encephalopathy (CTE)^50,51^. Our findings suggest that pathological phosphorylation patterns, but not phosphorylation generally, not only alters cooperation between tau molecules during envelope formation along microtubules, but also promotes the assembly of filaments relevant to tauopathies.

## Results

### Characterization of differentially phosphorylated recombinant human tau proteins

To generate differentially phosphorylated tau proteins, we expressed the longest isoform of human tau (441 amino acids, 2N4R) in bacteria (*E. coli*), insect cells (Spodoptera frugiperda 9 cells, Sf9) and mammalian cells (Human Embryonic Kidney Cells 293, HEK293), referred to as *E. coli* tau, Sf9 tau and HEK tau respectively hereafter. All proteins were purified via a N-terminal StrepII-superfolder GFP (sfGFP) tag and were analyzed by the sodium dodecyl sulfate-polyacrylamide gel electrophoresis (SDS-PAGE). Both Sf9 tau and HEK tau exhibited a slower migration rate compared with their non-phosphorylated cognate *E. coli* tau (Figure 1A), presumably due to the alteration of protein molecular weight or charge introduced by phosphorylation. The 2N4R tau protein has 85 potential phosphorylation sites, consisting of 45 serines, 35 threonines, and 5 tyrosines^3,52,53^. To identify the actual phosphorylated residues in our tau preparations, we used liquid chromatography-electrospray mass spectrometry (LC-MS). Our analysis revealed that the HEK tau protein contained 52 phosphorylated sites, while the Sf9 tau protein contained 58 phosphorylated sites detected. Both protein preparations share 38 common phosphorylation sites (Figure 1B). We conclude that human tau protein expressed in heterologous eukaryotic cells is differentially phosphorylated by endogenous cellular kinases.

**Figure 1.**
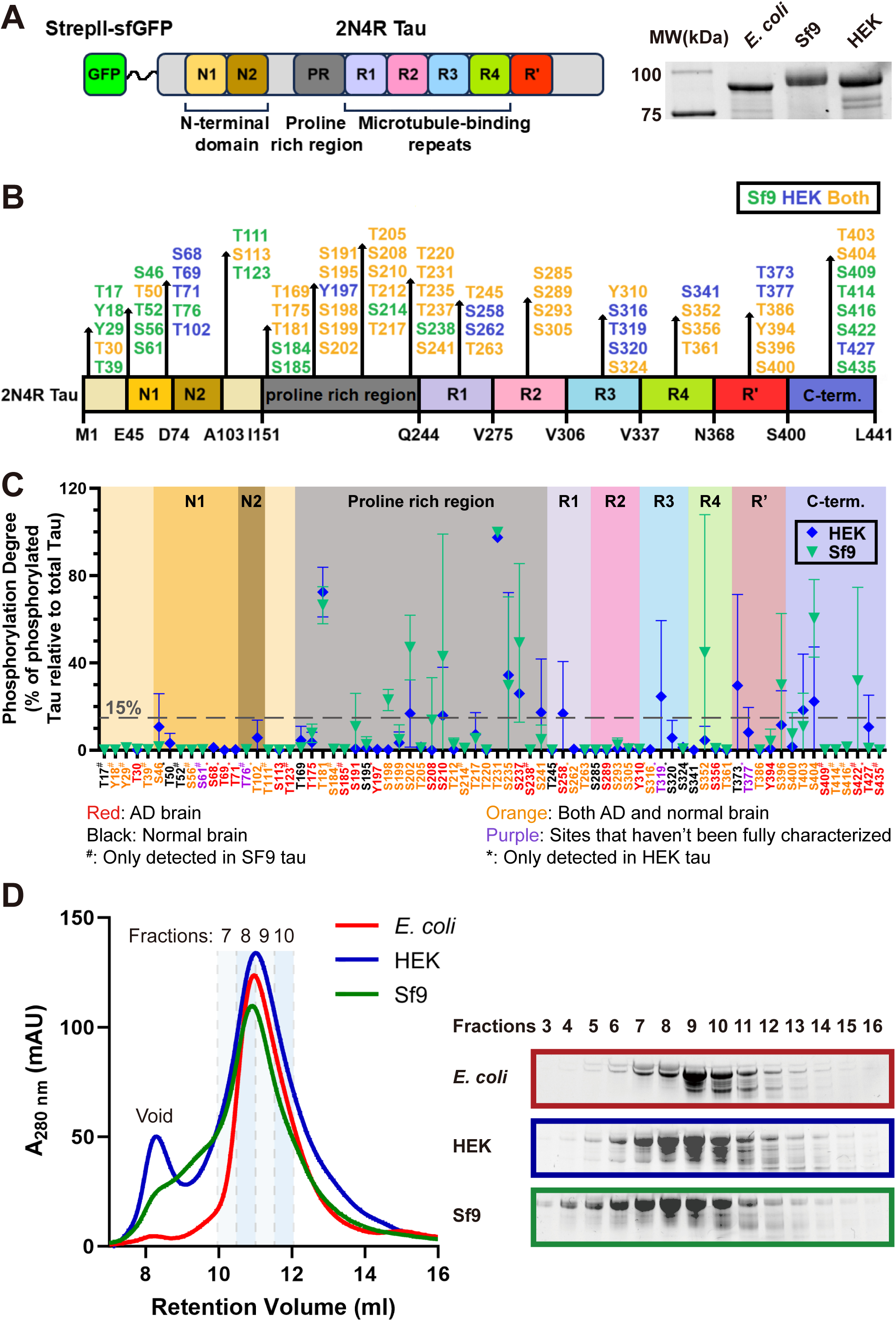
Characterization of differentially phosphorylated tau proteins. **A.** Left: Schematic of the human 2N4R tau construct used in this study, which contains N-terminal N1 and N2 insert, a proline rich region, 4 microtubule binding repeats and a C-terminal pseudo-repeat region (R’). Right: SDS–PAGE analysis of the differentially phosphorylated human 2N4R tau proteins which were expressed and were purified from different expression systems. Note: the precise definition of the microtubule binding repeats has not always been consistent in previous literature, we defined the microtubule binding repeats as residues 242-367, similar to a previous cryo-EM study, in which the R’ is not included^70^. **B.** Schematic showing the distribution of the phosphorylation sites in HEK tau and Sf9 tau detected by the mass spectrometry. Sites only detected in HEK tau are labeled dark blue, sites only detected in Sf9 tau are labeled dark green, sites detected in both HEK tau and Sf9 tau are labeled amber/yellow. **C.** Schematic showing the site-specific phosphorylation degrees at corresponding sites in the HEK tau (dark bule) and the Sf9 (dark green) tau based on the calculation from mass spectrometry. *N* = 2, Mean ± SD is shown. Annotation at the bottom shows the phosphorylation sites detected in either AD or normal brain (summarized from previous studies^59–65^). **D.** Size exclusion chromatograph of differentially phosphorylated tau proteins. Left: Profiles of the differentially phosphorylated tau proteins. Right: SDS-PAGE gel of peak fractions shows that there is only subtle differences between tau proteins purified from different expression systems.

To quantify the phosphorylation status of these tau proteins, we determined the site-specific phosphorylation levels of both HEK tau and Sf9 tau by comparing the signal intensity of phosphorylated peptides at specific sites to the total signal intensity of both phosphorylated and non-phosphorylated peptides within the same site^54,55^. This analysis revealed that the phosphorylation occupancy in most sites of both HEK tau and Sf9 tau was lower than ∼15% (Supplementary table 1). Most sites where the phosphorylation occupancy was higher than 15% were located outside of the microtubule binding repeat (Figure 1C). Specifically, in HEK tau, 7 sites located in the proline rich region (T181, S202, S210, T231, S235, S237 and S241), 3 sites located in the C-terminus (T373, T403 and S404) and 2 sites (S258 and T319) located in the microtubule binding repeat had an occupancy of 15% or higher (Figure 1C and Supplementary table 1). In the Sf9 tau protein, we identified 11 sites with phosphorylation occupancy greater than 15%. These sites are distributed across three regions: the proline-rich region (7 sites: T181, S198, S202, S210, T231, S235 and S237), the C-terminal region (3 sites: S396, S404, and S422), and the microtubule binding repeat (1 site: S352, Figure 1C and Supplementary Table 1). The distribution of phosphorylated sites in our Sf9 tau is similar to a previous report^56^. These results reveal that specific sites within the tau molecule are more susceptible to phosphorylation by endogenous kinases.

Among the highly phosphorylated sites, T181 and T231 are phosphorylated in both HEK tau and Sf9 tau, and have been previously reported as key sites in Alzheimer’s disease (AD)^57^. Phosphorylated T181 and T231 have been identified as biomarkers for AD^58^, and previous studies have shown that they are highly phosphorylated in Sf9 tau as well^49,56^. Additionally, phosphorylation at S241, S258, T319, and T373 is specific to HEK tau or is significantly higher in HEK tau than in Sf9 tau. Conversely, phosphorylation at S198, S202, S210, S237, S352, S396, S404 and S422 is specific to Sf9 tau or is significantly higher in Sf9 tau than in HEK tau (Figure 1C and Supplementary Table 1). Notably, phosphorylation at S210, S237, S258, and S422 has been only detected in AD patient brains (Figure 1C)^59–65^.

Previous research has shown that tau protein is typically phosphorylated at 2-3 sites in healthy brains, whereas in NFTs, it is aberrantly hyperphosphorylated at 5-9 sites^66,67^. Our findings are consistent with these reports, as we observed an average phosphorylation degree of approximately 9% (∼5 phosphorylated sites per molecule) in HEK tau, which is similar to the average phosphorylation degree of approximately 11% (∼6 phosphorylated sites per molecule) in Sf9 tau (Supplementary Table 1). We note that previous studies used the term “hyperphosphorylation” to describe the phosphorylation status of tau protein expressed in insect cells. However, this term is ambiguous, as our results show that the average phosphorylation degree of our proteins is only around 10% (Supplementary Table 1). To avoid confusion, we will use the term “phosphorylation” instead of “hyperphosphorylation” to describe the phosphorylation status of our tau proteins in the remainder of this study. These results demonstrate that while HEK tau and Sf9 tau share similar levels of phosphorylation, the phosphorylation patterns in these two proteins are distinct.

Given the distinct phosphorylation patterns observed in HEK tau and Sf9 tau, we sought to investigate whether these differences impact the conformation or ensemble oligomeric state of tau molecules in solution. To address these questions, we first employed size exclusion chromatography to analyze the apparent molecular size of both HEK tau and Sf9 tau, using non-phosphorylated *E. coli* tau as a reference control. The chromatographic profiles revealed that HEK tau eluted highly similarly to *E. coli* tau, whereas Sf9 tau eluted slightly earlier than both *E. coli* tau and HEK tau (Figure 1D). This suggests that the oligomeric state of these three tau proteins is similar under these conditions, with the slightly earlier elution of Sf9 tau potentially indicating a conformational change or a population of molecules in an altered oligomeric state. However, the limited resolution of this method makes it challenging to definitively distinguish between these possibilities.

Combined, these findings demonstrate that we have produced differentially phosphorylated tau proteins, and that these proteins exhibited a similar distribution of their phosphorylation sites but with unique phosphorylation patterns. Further, phosphorylation of tau does not dramatically alter its ensemble oligomeric state or hydrodynamic radius.

### Effects of phosphorylation on single molecule and ensemble microtubule binding behaviors of tau

Tau phosphorylation may influence its interaction with microtubules by altering the protein’s charge status and/or conformation, which could disrupt the formation of salt bridges between tau and microtubules^30–33,53,68–70^. To investigate whether phosphorylation affects the dynamics of tau molecules on microtubules, we first examined the behavior of differentially phosphorylated tau proteins at the single-molecule level. We tracked the binding dynamics of single SNAP-tagged 2N4R tau molecules along surface-immobilized taxol-stabilized microtubules at single molecule concentrations (Figure 2A). We quantified the single molecule dwell times and landing rates, revealing that phosphorylated tau molecules displayed an ∼ 2.5-fold reduced dwell time on microtubules (*E. coli* tau: ∼ 4.3 seconds, HEK and Sf9 tau: ∼ 1.75 seconds, (Figure 2B). Additionally, the landing rates of phosphorylated tau molecules were reduced by ∼ 2-10-fold (*E. coli* tau: 0.67 ± 0.43 nM^-1^μm^-1^s^-1^, HEK tau: 0.40 ± 0.28 nM^-1^μm^-1^s^-1^, Sf9 tau: 0.08 ± 0.05 nM^-1^μm^-1^s^-1^, (Figure 2C). These results indicate that tau phosphorylation decreases the initial binding rate and dwell time of tau with microtubules, consistent with previous bulk biochemical studies which showed the tau phosphorylation lowers its overall microtubule affinity^30–33^.

**Figure 2.**
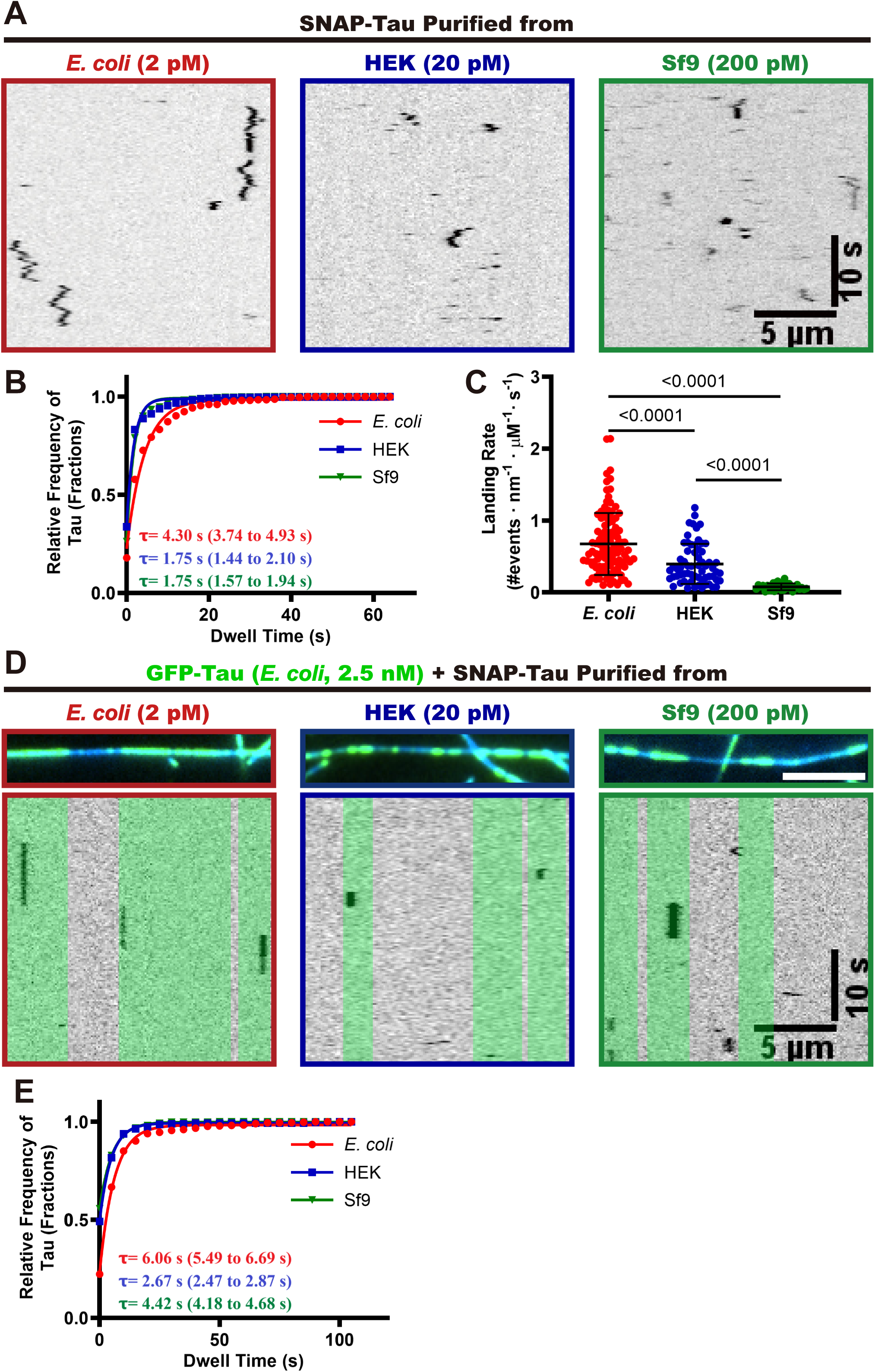
Single molecule analysis of the effects of phosphorylation on the tau-microtubule interaction. **A.** Kymographs of differentially phosphorylated single SNAP-tau molecules diffusing on bare microtubules. Scale bars: 10 sec. and 5 µm. **B.** Quantification of the dwell times of differentially phosphorylated tau proteins on microtubules. Cumulative frequency of the dwell time is plotted for each population of tau proteins and fit to a one-phase exponential decay function. The characteristic dwell times derived from the fits (τ) and the corresponding 95% confidence intervals (95% CI, in parentheses) are indicated. Events longer than three pixels were selected and quantified and two independent experiments were performed for each condition. *E. coli* tau: n = 330, R^2^ = 0.978. HEK tau: n = 831, R^2^ = 0.973. Sf9 tau: n = 726, R^2^ = 0.990. *N* = 2, Mean ± SD is shown. **C.** Quantification of landing rates of differentially phosphorylated tau proteins on microtubules. For quantification, microtubules were quantified from two independent experiments for each tau protein. *E. coli* tau: n = 106 microtubules quantified. HEK tau: n = 63. Sf9 tau: n = 31 microtubules quantified. *P* values are calculated from an unpaired, two-tailed *t*-test. *N* = 2, Mean ± SD is shown. **D.** Kymographs of differentially phosphorylated SNAP-tau proteins within non-phosphorylated GFP-tau envelopes on microtubules. Tau envelope boundaries are colored green on the kymographs. Scale bars: 10 sec. and 5 µm. **E.** Quantification of the dwell times of differentially phosphorylated SNAP-tau proteins within envelopes formed by non-phosphorylated GFP-tau. Cumulative frequency of the dwell time is plotted for each population of tau proteins and fit to a one-phase exponential decay function. The characteristic dwell times derived from the fits (τ) and the corresponding 95% CI (in parentheses) are indicated in the bottom left corners. *E. coli* tau: n = 430, R^2^ = 0.993. HEK tau: n = 352, R^2^ = 0.998. Sf9 tau: n = 353, R^2^ = 0.998.

We and others have proposed that one of the key functions of the tau protein is its ability to form cohesive tau envelopes on the microtubule surface, which acts as a selectively permeable barrier to regulate the activities of other microtubule-associated proteins (MAPs)^13,14^. Within this envelope, tau proteins exhibit cooperative interactions with each other, resulting in longer dwell times compared to tau proteins outside the envelope^13,14^. To investigate whether phosphorylation affects the interaction of tau proteins within these envelopes, we employed a single-molecule assay to measure the dwell times of differentially phosphorylated tau proteins within envelopes formed by non-phosphorylated *E. coli* tau. In this assay, 2.5 nM of GFP-tagged *E. coli* tau were preincubated with taxol-stablized microtubules to form envelopes, followed by the introduction of differentially phosphorylated and SNAP-labelled tau proteins. Our results show that all three tau proteins can engage with the envelopes formed by non-phosphorylated *E. coli* tau (Figure 2D). We next examined the dwell times of the tau proteins within the *E. coli* tau envelopes. The measured dwell time of single *E. coli* tau molecules was ∼ 6.06 seconds, while the dwell times of phosphorylated HEK tau and Sf9 tau molecules was reduced to ∼ 2. 7 seconds and ∼ 4.4 seconds, respectively (Figure 2D and 2E). Notably, all three tau proteins exhibited increased dwell times within the envelopes compared to their dwell times outside of the envelopes (Figure 2B), confirming the existence of cooperative interactions between the differentially phosphorylated tau proteins within the envelopes. The shorter dwell times of phosphorylated tau molecules within tau envelopes suggests that phosphorylation impairs the cooperative interaction between phosphorylated and non-phosphorylated tau proteins within the envelopes, consistent with a previous study which found that Cdk5 phosphorylation can induce disassembly of envelopes formed by non-phosphorylated bacterial tau^58^. Interestingly, we found that HEK tau exhibited the shortest dwell times within the envelopes, ∼ 2.3 times shorter than *E. coli* tau. We speculate this may be attributed to the additional phosphorylation sites (T373 and T377) in the pseudo-repeat Region (R’) of HEK tau (Figure 1C)), which has been reported to regulate the cooperative interaction between tau molecules on microtubules^13^. Our results show that phosphorylation impairs the cooperative interaction between tau molecules within envelopes formed by non-phosphorylated tau, and that specific phosphorylation sites may differentially impact these interactions that underly tau envelope formation.

### Phosphorylated tau retains the capability to form envelopes and to compact the microtubule lattice

We have shown that phosphorylation reduces the affinity of single tau molecules for microtubules and diminishes the cooperative interactions within envelopes formed by non-phosphorylated tau. We next sought to investigate whether phosphorylation abolishes tau’s ability to form envelopes. To address this question, we examined the capacity of differentially phosphorylated tau proteins to form envelopes on microtubules at various concentrations. Our results show that all three tau proteins retain the ability to form envelopes on microtubules, although Sf9 tau and HEK tau require substantially higher concentrations to do so (Figure 3A). This observation is consistent with previous reports that higher concentrations of Sf9 tau are required to form envelopes on microtubules^13,56^. These findings indicate that while phosphorylation diminishes tau’s microtubule affinity and impairs the cooperative interactions within envelopes formed by non-phosphorylated tau, it does not fully eliminate its ability to form envelopes on microtubules.

**Figure 3.**
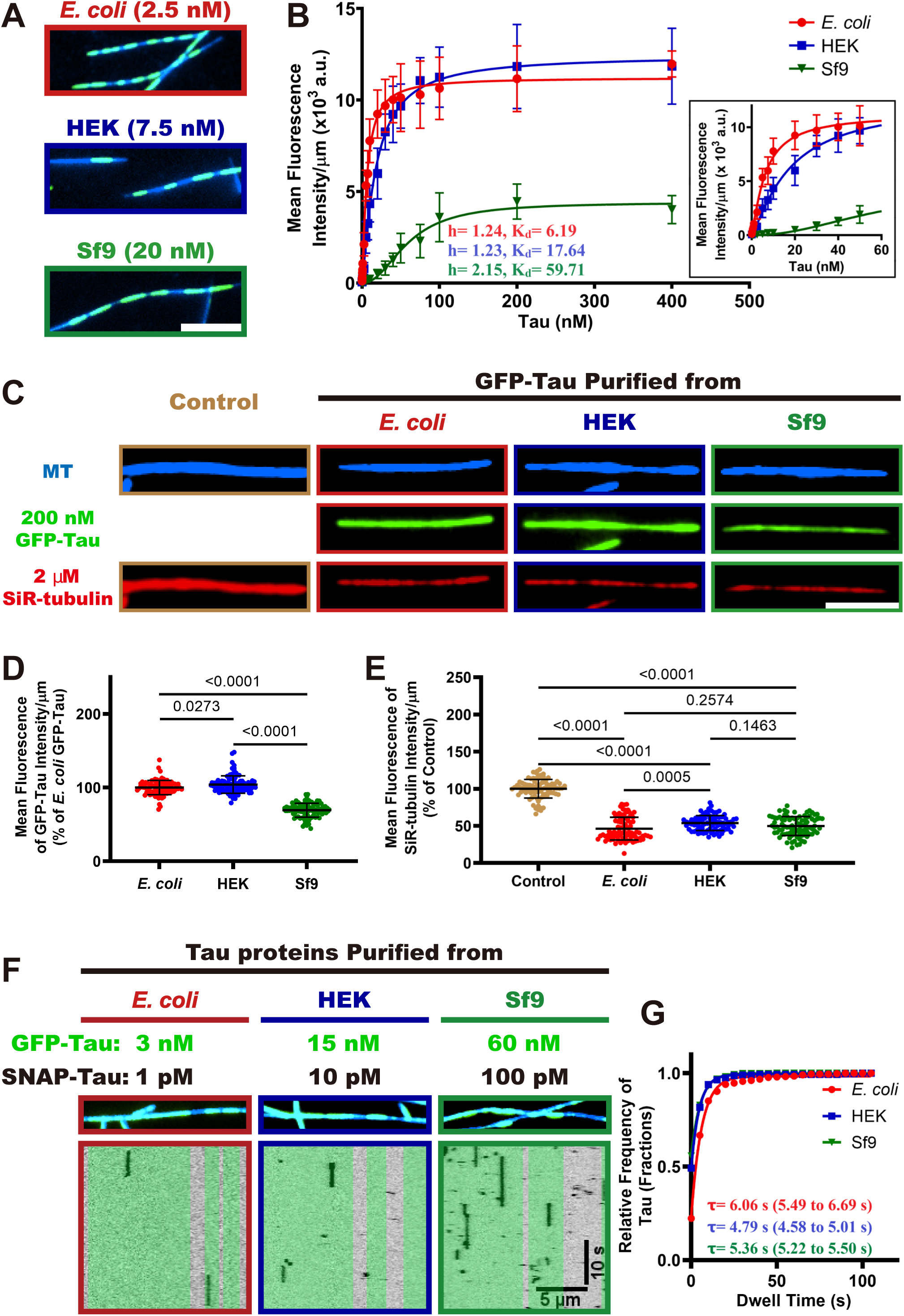
Specific phosphorylation patterns alter cooperative interactions between tau proteins on microtubules. **A.** TIRF images showing GFP-tau envelopes on taxol-stabilized microtubules. Scale bar: 5 µm. **B.** Quantification of differentially phosphorylated tau proteins binding to microtubules across a range of tau concentrations. Binding curves were fitted to the Hill equation to derive the Hill coefficient and *K_d_*. For each concentration of corresponding tau protein, *N* = 2, n = 60 microtubules. Mean ± SD is shown. Hill coefficients and *K_d_* are indicated. 95% CI for Hill coefficients: 1.16 to 1.33 (*E.coli* tau), 1.17 to 1.30 (HEK tau), 1.97 to 2.34 (Sf9 tau). 95% CI for *K_d_*: 5.84 to 6.60 (*E.coli* tau), 16.66 to 18.70 (HEK tau), 56.93 to 62.75 (Sf9 tau). Insert showing the initial time course of binding. **C.** TIRF images showing microtubules and differentially phosphorylated tau proteins at saturating concentration (200 nM), along with SiR-tubulin bound along the microtubules. Scale bar: 5 µm. **D.** Quantification of the average fluorescence intensities of differentially phosphorylated GFP-tau proteins on the microtubules. *E. coli* tau: n = 90. HEK tau: n = 90. Sf9 tau: n = 90. *P* values are calculated from an unpaired, two-tailed *t*-test. *N* = 2, Mean ± SD is shown. **E.** Quantification of the average fluorescence intensities of SiR-tubulin along the microtubules. Control condition: n = 90 microtubules. *E. coli*: n = 90. HEK tau: n = 90. Sf9 tau: n = 90. *P* values are calculated from an unpaired, two-tailed *t*-test. *N* = 2, Mean ± SD is shown. **F.** Kymographs of differentially phosphorylated SNAP-tau proteins within envelopes formed by corresponding GFP-tau proteins on the microtubules. Tau envelope boundaries are colored green on the kymographs. Scale bars: 10 sec and 5 µm. **G.** Quantification of the dwell times of differentially phosphorylated tau proteins within envelopes on the microtubules. Cumulative frequency of the dwell time is plotted for each population of tau proteins and fit to a one-phase exponential decay function. The characteristic dwell times derived from the fits (τ) and the corresponding 95% CI (in parentheses) are indicated. (the *E. coli* tau data were re-plotted from figure 2E for comparison). *E. coli* tau: n = 413, R^2^ = 0.993. HEK tau: n = 311, R^2^ = 0.999. Sf9 tau: n = 430, R^2^ = 0.9995.

Because tau envelope formation is a cooperative process^17^, we next sought to investigate the impact of phosphorylation on the cooperative interaction between tau molecules during envelope formation on microtubules. We measured the average fluorescence intensity of differentially phosphorylated tau proteins at various concentrations, and the data were fitted to the Hill equation (Figure 3B). We measured very similar Hill coefficients of 1.24 (95% Confidence Intervals, 1.16 to 1.33) for *E. coli* tau, and 1.23 (95% CI, 1.17 to 1.30) for HEK tau, indicating that the phosphorylation pattern found on HEK tau does not dramatically alter the cooperative interaction between HEK tau molecules during envelope formation (Figure 3B). In contrast, the Hill coefficient of Sf9 tau was elevated to 2.15 (95% CI, 1.97 to 2.34), suggesting that this phosphorylation pattern, or phosphorylation of specific sites, enhances the cooperative interaction between Sf9 tau molecules. Furthermore, the dissociation constant (K_d_) values of *E. coli* tau, HEK tau, and Sf9 tau were 6.19 nM, 17.64 nM, and 59.71 nM respectively (Figure 3B), confirming that phosphorylation decreases the microtubule affinity of tau protein, consistent with our single-molecule results (Figure 2). These findings collectively demonstrate that phosphorylation does not strongly disrupt, and can even enhance, the cooperative interaction between tau proteins within envelopes, while also reducing their overall microtubule affinity. Notably, the maximum binding capacity (B_max_) of Sf9 tau was ∼ 3-fold lower than that of *E. coli* and HEK tau (Figure 3B), which may be attributed to the lower landing rate of Sf9 tau (Figure 2A-C). We speculate that Sf9 phosphorylation may also induce a conformational change, as supported by our gel filtration results (Figure 1D), which may conceivably affect envelope formation and density. Further investigation is needed to confirm this hypothesis

We have proposed that envelopes play a crucial role in regulating the accessibility of microtubule-associated proteins (MAPs) to the microtubule lattice^13,14^. One mechanism for this activity may be through compaction of the underlying tubulin lattice^17,18^. Given that the B_max_ of Sf9 tau is significantly lower than that of *E. coli* tau and HEK tau, we investigated whether Sf9 tau was competent to induce microtubule lattice compaction. We employed a SiR-tubulin assay to investigate the ability of differentially phosphorylated GFP-tau proteins to compact microtubule lattices, using taxol-stabilized microtubules with expanded tubulin lattices and 2 μM SiR-tubulin preincubated on a coverslip. We then added the phosphorylated tau proteins at saturating concentrations (200 nM) and measured the fluorescence intensities of both GFP-tau and SiR-tubulin using TIRF microscopy. Our results show that the saturating fluorescence intensity of Sf9 tau is approximately 69% lower than *E. coli* tau (100%), while HEK tau was very similar to *E. coli* tau (104%, Figure 3C-D). This is consistent with the binding curve data obtained at saturating concentrations (Figure 3B). The addition of *E. coli* tau to the microtubules resulted in a significant decrease in SiR-tubulin fluorescence intensity by ∼ 50% (Figure 3C, E), indicating that *E. coli* tau compacts the microtubule lattice and competes with SiR-tubulin for binding, consistent with prior results^17^. Interestingly, the SiR-tubulin intensity in the presence of HEK tau and Sf9 tau were similar to those of *E. coli* tau, suggesting that these differentially phosphorylated tau proteins exhibit similar capabilities to compact the microtubule lattice (Figure 3C, E) at saturating concentrations^17^.

We next measured the dwell times of single tau molecules within envelopes formed from the same type of tau preparation (non-phosphorylated or phosphorylated). Non-phosphorylated *E. coli* tau showed the longest dwell times, while HEK cell tau molecules showed somewhat extended dwell times inside phosphorylated HEK tau envelopes as compared to within dephosphorylated *E. coli* tau envelopes (Figures 3F-G and 2D-E). This result indicates increased interactions between phosphorylated HEK cell tau molecules within envelopes. Phosphorylated Sf9 cell tau molecules also showed modestly enhanced dwell times within Sf9 cell tau envelopes as compared to within *E. coli* tau envelopes (Figures 3F-G and 2D-E). These results further confirm our observations that tau phosphorylation while reducing overall microtubule affinity, also impacts tau’s cooperative interactions (Figure 2B). We conclude that phosphorylated tau retains the capability to form cooperative envelopes and compact the microtubule lattice, which in turn regulates the microtubule-based activities, consistent with recent results showing that phosphorylated tau partially retains its function which works as a selectively permeable barrier^56^.

### Phosphorylated tau forms small oligomers in solution

Hyperphosphorylation of tau has been implicated in its oligomerization and the pathogenesis of neurodegenerative tauopathies^20–24^. Previous studies have demonstrated that tau hyperphosphorylation in Sf9 and HEK cells enhances oligomerization^71,72^. To more precisely characterize the oligomeric state of differentially phosphorylated tau proteins, we employed mass photometry across a range of nanomolar tau concentrations. Recombinantly expressed tau from *E. coli* remained predominantly monomeric (∼ 95%) across a 25-fold concentration range, with no detectable dimer formation (Figure 4). Both HEK- and Sf9-derived tau existed primarily as monomers at 1 nM, with no observable dimers. However, increasing the concentration to 5 nM promoted oligomerization, resulting in 22% and 11% dimer formation for HEK and Sf9 tau, respectively (Figure 4). At 25 nM, dimeric species increased to 32% and 12% for HEK and Sf9 tau, respectively, with HEK tau also exhibiting 4% trimer formation (Figure 4). These findings corroborate previous biochemical studies demonstrating increased oligomerization propensity of Sf9-expressed phosphorylated tau^71^ and highlight the increased tendency of HEK-derived tau to form higher-order soluble oligomers at elevated concentrations. Differential phosphorylation patterns likely contribute to the observed variations in oligomerization behavior between HEK and Sf9 tau and require further investigation. These results demonstrate that tau phosphorylation favors the formation of small soluble tau oligomers starting with dimers and trimers.

**Figure 4.**
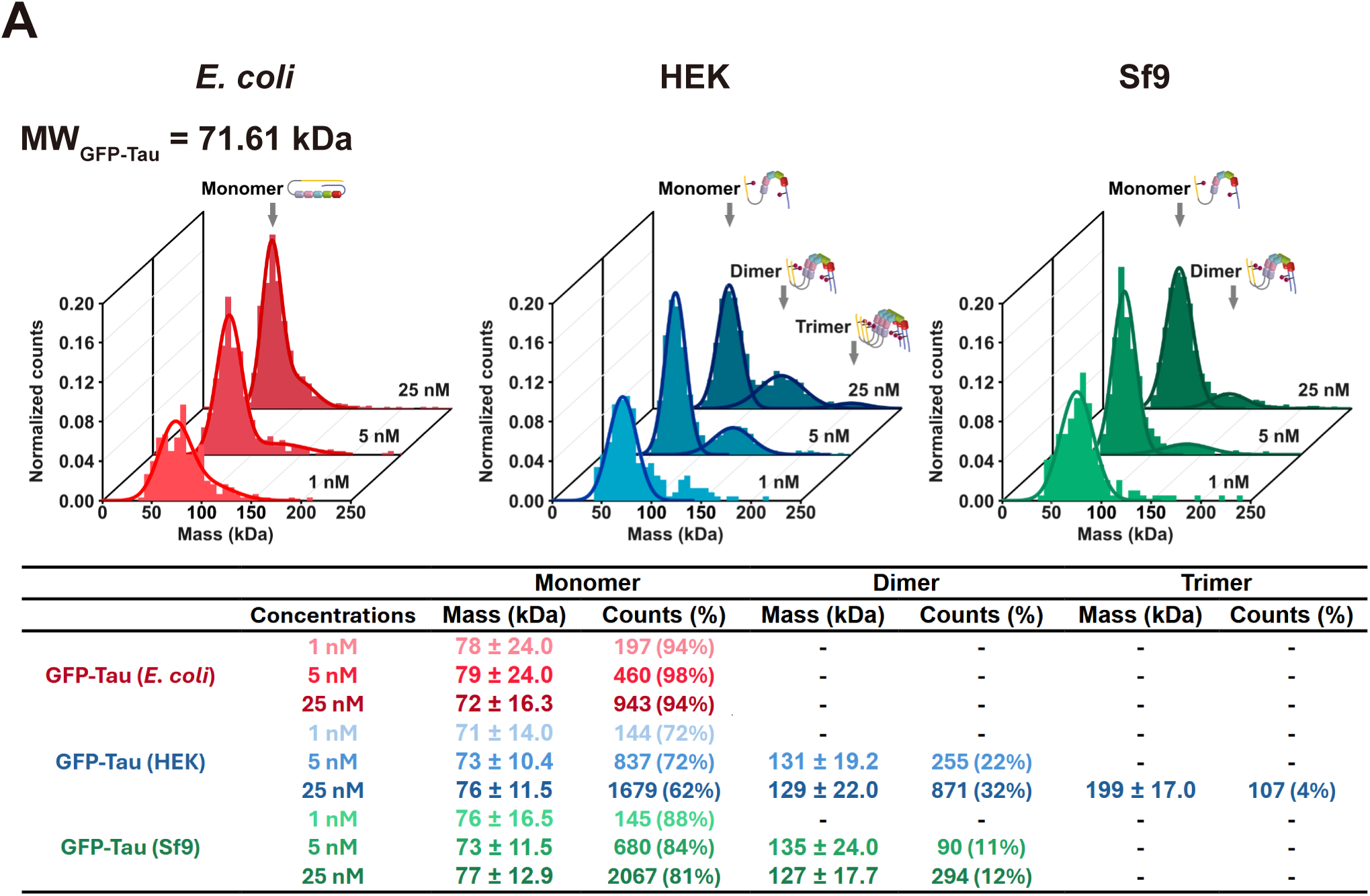
Phosphorylation promotes oligomerization of tau in solution. Characterization of the oligomeric state of differentially phosphorylated tau proteins by mass photometry. Histograms in each panel show the normalized particle counts (y-axis) of differentially phosphorylated tau proteins at the indicated molecular mass (x-axis). For each protein, the molecular mass was measured at three different indicated concentrations (indicated in z-axis). Dark lines indicate Gaussian fits to the peaks. The calculated mass, percentage of particles, and total particle counts of each oligomeric state are indicated. Cartoons indicate the hypothesized oligomeric state of each indicated peak. The theoretic molecular weight of GFP-tau was indicated in the top left.

### Pathological tau phosphorylation pattern facilitates the assembly of full-length tau into CTE-like NFT fibrils

A common feature of tauopathies is the intracellular accumulation of NFTs composed of hyperphosphorylated tau^20–25^. While hyperphosphorylated tau is a major component of these filaments, the causal relationship between tau phosphorylation and NFT assembly remains a subject of ongoing debate^39^. Recent studies have achieved the *in vitro* assembly of tauopathy-relevant filaments without the need for polymerization-promoting cofactors^50,51,73,74^. To investigate the role of tau phosphorylation in filament assembly, we conducted an *in vitro* polymerization assay to generate tau fibrils that adopt the same conformation as those isolated from tauopathy disease brains. In this assay, we used a tau fragment derived from the AD/CTE tau filament core, spanning residues 297-391 of the protein^26,28^. This fragment, known as dGAE^75^, encompasses intact regions of the microtubule binding repeat R3 and R4, as well as part of R2 and the pseudo-repeat (Figure 5A). Previous studies have demonstrated the successful polymerization of dGAE into AD-like and CTE-like filaments *in vitro*, in the absence of aggregation-inducing cofactors, which exhibit identical atomic structures to those isolated from AD and CTE brains^50,51^.

**Figure 5.**
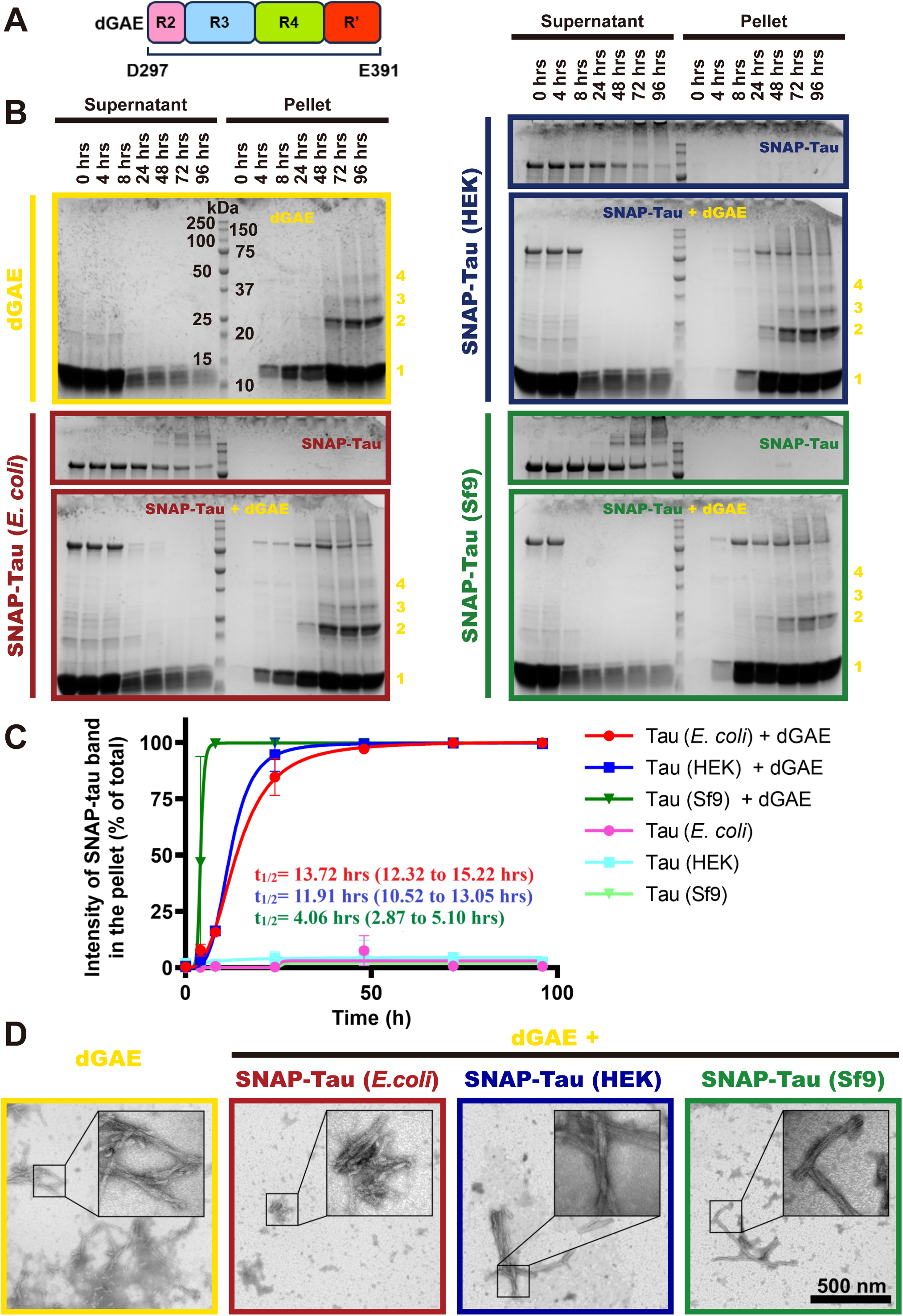
Specific phosphorylation patterns promote tau co-assembly into CTE-like fibrils formed by the tau fragment dGAE. **A.** Diagram of dGAE, which contains a C-terminal portion of R2, intact R3, R4 and N-terminal portion of R’. **B.** Coomassie-stained SDS-PAGE gels showing the assembly kinetics of CTE-like fibrils indicated by pelleting via centrifugation. dGAE bands representing inferred oligomeric states are indicated by numbers indicating those states (1: monomer, 2: dimer, 3: trimer, etc.). **C.** Quantification of the co-assembly kinetics of the differentially phosphorylated tau proteins. The fractions of the differentially phosphorylated full-length tau proteins in the pellet at the indicated time points in each condition were evaluated by comparing intensities of full-length tau in the pellet versus in total (pellet + supernatant) in each condition. Curves were fitted to one-site-specific binding with Hill slope. The calculated *t_1/2_* and the corresponding 95% confidence intervals (95% CI, in parentheses) are indicated, *N* = 2, Mean ± SD is shown. **D.** Negative-stain transmission electron microscopy (TEM) images of filaments assembled from dGAE or co-assembled from dGAE + differentially phosphorylated full-length tau proteins. Scale bar: 500 nm.

We formed CTE-like filaments by continuous agitation of 400 μM dGAE in previously established buffer conditions^50,51^. Analysis by ultracentrifugation and SDS-PAGE revealed assembly initiation before 4 hours, detectable higher-order oligomers at 24 hours, and near-complete incorporation of dGAE into filaments by 96 hours (Figure 5B). Negative-stain transmission electron microscopy (TEM) confirmed the characteristic twisted filament structure consistent with previous reports of CTE-like filaments (Figure 5D)^50,51^.

To assess the impact of phosphorylation on incorporation of tau molecules into the growing filaments, differentially phosphorylated full-length SNAP-tagged tau proteins were co-assembled with dGAE. A 1:99 molar ratio of SNAP-tau (4 μM) to dGAE (396 μM) was used to minimize any steric effects from the SNAP tag^76^. Parallel assays with SNAP-tau alone (4 μM) served as controls. Ultracentrifugation and SDS-PAGE analysis revealed near-complete incorporation of all differentially phosphorylated SNAP-tau species into the pellet fraction after 96 hours of co-assembly, whereas in the absence of dGAE, full-length tau remained in the supernatant (Figure 5B-C). Negative-stain TEM confirmed the formation of twisted filaments in all co-assembly samples after 96 hours (Figure 5D), indicating incorporation of the differentially phosphorylated tau into CTE-like filaments in the presence of dGAE, while full-length tau alone did not assemble under these conditions.

To quantify the impact of phosphorylation on aggregation kinetics, the fraction of pelleted full-length tau was determined at each time point and fitted using the Hill equation (Figure 5C). The t_1/2_ values for *E. coli* tau and HEK tau were similar (∼ 14 hours and ∼ 12 hours, respectively). However, the t_1/2_ for Sf9 tau was significantly faster (∼ 4 hours) (Figure 5B), demonstrating that the Sf9 tau phosphorylation pattern strongly promotes co-assembly into CTE-like filaments. While both HEK and Sf9 tau exhibit similar overall phosphorylation levels (∼ 5-6 sites per molecule), their phosphorylation patterns differ. Given the reported similarity between the phosphorylation pattern of insect cell-expressed tau and AD-derived tau^49^, Sf9 tau appears to be a robust pathological tau model system. Our co-assembly assay reveals that particular phosphorylation patterns facilitate the assembly of full-length tau molecules into pathologically relevant fibril structures.

## Discussion

Tau phosphorylation is a critical regulator of both physiological and pathological processes in mammals. Physiologically, tau phosphorylation is implicated in diverse functions, including development^44–46^, hibernation^41–43^, normal sleep^77^, and anesthesia-induced modulation of consciousness^78,79^. During these processes, transient increases in tau phosphorylation at multiple sites are reported, which subsequently return to lower physiological levels^41–43,47,77^. Conversely, pathologically altered tau phosphorylation disrupts microtubule (MT) binding affinity and induces conformational changes, potentially leading to tau mislocalization from axons to the cytosol^40^ or promoting aggregation^80^. Consistent with this, highly phosphorylated tau is a major component of paired helical filaments (PHFs) in several tauopathies^20,24,25^. Intriguingly, while physiological tau phosphorylation appears reversible, that observed in pathological conditions appears to be irreversible, despite similarities in phosphorylation levels between fetal and Alzheimer’s disease (AD) tau^39,45,46,48^. Furthermore, the functional consequences of phosphorylated tau in physiological contexts and the precise timing of tau phosphorylation relative to insoluble aggregate formation in AD and other tauopathies remain unclear^39^.

To help elucidate the role of tau phosphorylation, we characterized the functional impacts of differentially phosphorylated tau proteins on microtubule binding and tau envelope formation at the single-molecule level. We also investigated the role of phosphorylation in tau oligomerization at nanomolar concentrations and its influence on tauopathy-related fibril assembly. We utilized two distinct forms of phosphorylated tau: HEK tau, derived from human embryonic kidney (HEK) 293 cells, representing a potentially physiological form, and Sf9 tau, previously shown to exhibit similar phosphorylation patterns to tau isolated from AD brains^49^, thus serving as a model for pathological tau.

Previous studies have suggested that the addition of phosphate groups can induce local charge effects, which in turn alter the conformation and molecular interactions of tau^81,82^. In our study, we observed that HEK tau and Sf9 tau exhibit similar phosphorylation levels, with approximately 5-6 phosphorylated sites per molecule. As a result, both tau preparations showed reduced affinity for MTs compared to non-phosphorylated tau (Figure 2), consistent with previous findings that specific phosphorylation at sites S214, T231, S235, or S262 inhibits MT interactions (Figure 1C)^32,34^ However, despite this similarity, Sf9 tau displayed a significantly lower microtubule affinity than HEK tau, which might be attributed to the higher phosphorylation levels at S237 and S352 in Sf9 tau (Figure 1C). Specifically, the phosphorylation at S237 is at least 2-fold higher than in HEK tau, which is noteworthy given that phosphorylation of S235, a nearby residue, has been shown to inhibit microtubule binding^19^. Additionally, S352 is located in a conserved 12-residue stretch within R4 of the microtubule binding repeats, the corresponding residue in R1 (S258), has been shown to form a hydrogen bond with α-tubulin E434^70^, perhaps hinting at a mechanism for phosphorylation-induced disruption of microtubule binding. The corresponding residue S352 in R4 may also form a hydrogen bond with α-tubulin E434, the phosphorylation of S352 may also disrupt the interaction between tau and tubulin. Furthermore, S352 is located within the microtubule binding repeat R4 (Figure 1C), which is proximal to Ser356 within the KXGS motif. The KXGS motifs are found in the microtubule binding repeats of tau, MAP2 and MAP4 (X = I or C), theses KXGS motifs are targets of the microtubule-affinity regulating kinase (MARK)^30,83–85^. Previous experiments have demonstrated that phosphorylation within these motifs reduce MAP affinity for microtubules^30,84–86^. These findings suggest that the distinct phosphorylation patterns, and therefore charge distributions, between HEK tau and Sf9 tau may contribute to their differing microtubule affinities.

Despite reduced MT affinity, both HEK and Sf9 tau retained the ability to form tau envelopes and compact the MT lattice *in vitr*o (Figure 3C, E), suggesting the preservation of this crucial tau function even in pathologically phosphorylated tau. This observation is consistent with previous data showing that both *E. coli* and Sf9 tau form envelopes along microtubules, albeit at different concentrations^13,56^. To compensate for its reduced affinity, we found that Sf9 tau exhibited increased cooperative interactions on MTs (Figure 3B). While the underlying mechanism remains unclear, we observed differential phosphorylation at sites T373, T377, and S396 within the Pseudo-Repeat Region (R’) (Figure 1C), which has been previously implicated in cooperative tau interactions^13^. Site-specific mutagenesis studies will be essential to further dissect these mechanisms.

We observed that both HEK and Sf9 tau displayed concentration-dependent oligomerization in solution at nanomolar concentrations (Figure 4), indicating that phosphorylation of tau favors the formation of small soluble tau oligomers in solution. Given the relatively high reported abundance of tau in the neuronal cytoplasm (∼ 2 μM)^39^ and the presence of 2-3 phosphorylation sites in healthy brains^66,67^, our findings suggest that small tau oligomers may be present in the cytoplasm under physiological conditions. Consistent with this, a recent study show that tau can form dimeric and trimeric complexes under physiological conditions in cultured neurons and under non-aggregated conditions in engineered BSC-1 and QBI-293 cell lines^87^.

In our fibril co-assembly assays, Sf9 tau more readily co-assembled with the dGAE to form CTE-like filaments than either HEK or *E. coli* tau (Figure 5B, C). Given the similar overall phosphorylation levels between Sf9 and HEK tau, our results suggest that the pattern of phosphorylation, rather than overall level, is a crucial determinant for tauopathy-relevant filament assembly. Specifically, enhanced phosphorylation at S210 and S422 in Sf9 tau may facilitate co-assembly, as these sites are more highly phosphorylated in Sf9 tau than HEK tau, and phosphorylation at both sites is associated with AD (Figure 1C)^61–65^. Furthermore, phosphorylation at S208, near S210, as well as S422, is known to promote aggregation^35,36,38^. Phosphorylation at S262, which inhibits *in vitro* aggregation of a truncated tau fragment that encompasses the microtubule-binding repeats (termed K18)^88^, was also detected in our HEK tau preparation (Figure 1C), likely contributing to its reduced co-assembly propensity. Further site-specific mutagenesis will be crucial to define the precise roles of these individual sites. Importantly, neither of the differentially phosphorylated tau proteins formed filaments independently at the low concentration conditions we tested. However, we find that phosphorylation promotes oligomerization of tau in solution at nanomolar concentrations (Figure 4), underscoring the potential for phosphorylation to play multiple roles in tau aggregation, both promoting oligomerization and influencing filament formation in possibly distinct ways. Deciphering the precise mechanisms of these processes will be the focus of future research.

Our mass photometry data demonstrate that phosphorylated tau proteins exhibit a heightened propensity to form small oligomers in solution. Furthermore, co-assembly assays revealed that phosphorylated Sf9 tau more readily incorporates into large fibrils when combined with dGAE. These findings raise an intriguing question: is there a mechanistic link between tau oligomerization and fibrillization, or do these soluble oligomers represent an initial step in the fibrillization pathway? Interestingly, our assays indicate that phosphorylated HEK tau, despite showing the highest oligomerization propensity in solution, did not display significantly different co-assembly properties compared to non-phosphorylated E. coli tau. This suggests that the small oligomers formed by phosphorylated tau may not necessarily serve as the nucleation point for tau fibrillization, consistent with a previous study in engineered cells^87^, which found that physiological tau oligomers (dimers and trimers) are distinct from pathological aggregates with respect to phosphorylation state. Our work further highlights this distinction by revealing that specific phosphorylation patterns dictate the susceptibility of tau to transition into the aggregated state.

Supporting this notion, a recent study demonstrated that dimeric and trimeric tau species exist in both non-aggregated and aggregated states in engineered cell lines, adopting distinct conformations in each state^87^. Given that HEK tau and Sf9 tau in our study likely adopt different conformations in solution (Figures 1 and 4), we propose that molecular conformation, rather than the overall phosphorylation status, is a critical determinant in tau filament assembly. This highlights the importance of conformational dynamics in regulating tau aggregation and fibrillization pathways.

Recent studies have shown that liquid–liquid phase separation (LLPS) plays a crucial role in the formation of tau aggregations and can induce pathological tau conformations in solution (also known as tau liquid droplets), the LLPS is driven by multivalent interactions between tau molecules^89–91^. Moreover, our previous study revealed that on the microtubule surface, cooperative tau interactions that lead to envelope formation share some properties with solution-based LLPS^13^. Additionally, several recent studies show that phosphorylation drives tau LLPS and promotes droplets formation both *in vitro* and in cells^90,92^. Interestingly, the phosphorylated tau produced from Sf9 cells displayed fastest droplet formation rate and biggest droplet size among these differentially phosphorylate tau proteins tested^90^. These results hint that pathological pattern tau phosphorylation facilitated the interaction of tau molecules in solution via LLPS, consistent with our results on microtubules and in aggregation assays. Thus, we speculate that phosphorylation-impacted LLPS could be a mechanism for enhanced tau interactions leading to potential conversion into the pathological state.

In summary, our study revealed that the pathological pattern tau phosphorylation facilitates the cooperative interaction of tau molecules along microtubules potentially as a way to compensate for reduced microtubule affinity. Additionally, the pathological phosphorylation pattern also promotes tau molecules to co-assemble with dGAE into CTE-like filaments (Figure 6). Our study provides novel insights into the role of tau phosphorylation in regulating tau microtubule interactions and envelope formation, oligomerization, and tauopathy-related filament assembly. We find that the pattern of phosphorylation, but not the overall level of phosphorylation, is a key determinant in the conversion of tau from a highly soluble unstructured molecule to the pathologically folded filament structure observed in tauopathy disease.

**Figure 6.**
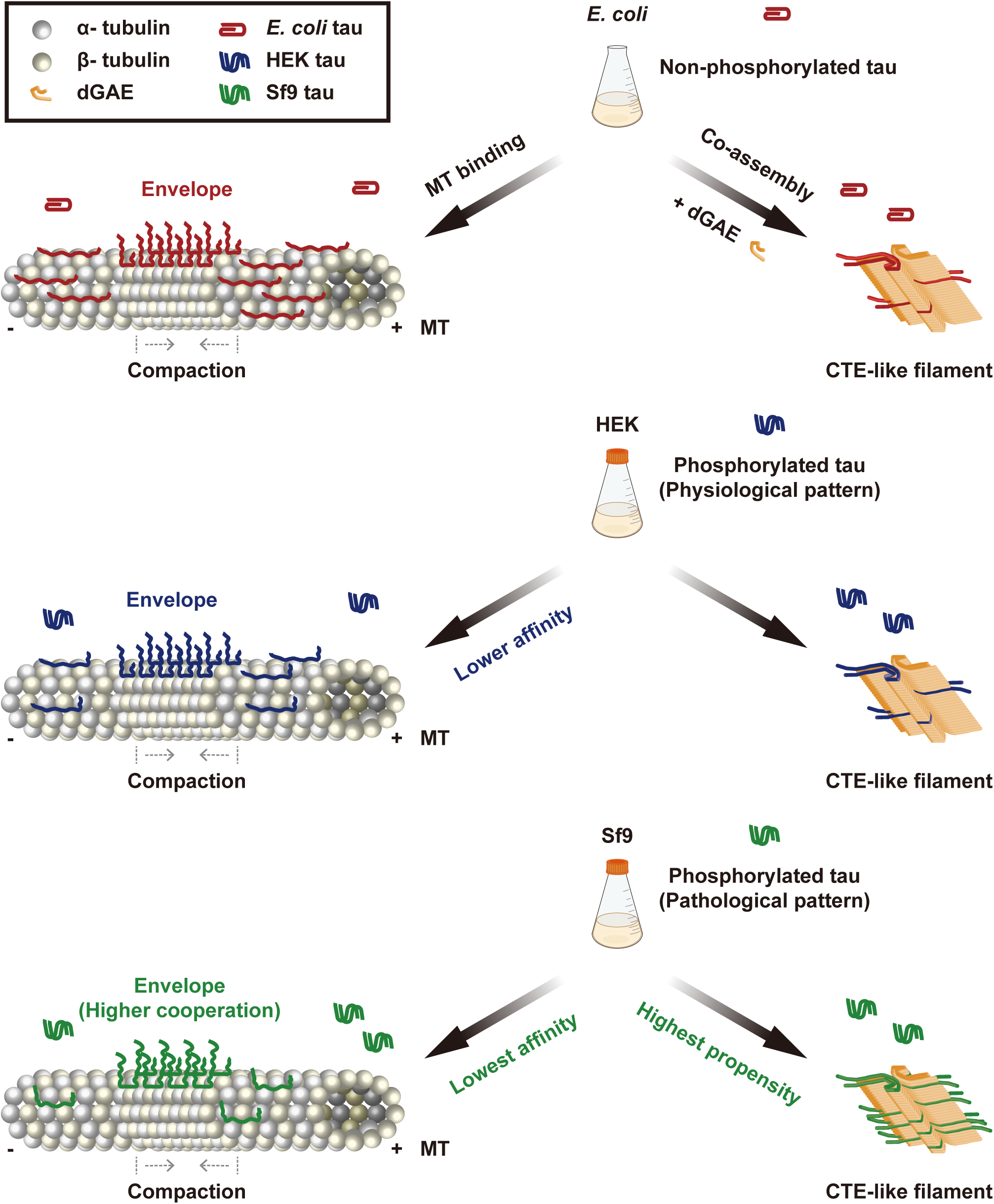
Summary of effects of tau phosphorylation on microtubule interactions and the tauopathy fibril co-assembly. Schematic showing a model for the effect of tau phosphorylation on microtubule interactions and CTE-like filament co-assembly. Middle panels showing the recombinant human 2N4R tau proteins were expressed and purified from different cellular sources: *E.coli* tau represent non-phosphorylated protein, HEK tau represents a physiological-like phosphorylation pattern, and Sf9 tau represents a pathological-like phosphorylation pattern. Left panels showing the interactions between the microtubule and differentially phosphorylated tau proteins. Although phosphorylation impairs the overall microtubule affinities of tau, all three types of tau proteins can form cooperative envelopes along microtubules and retain the ability to compact the microtubule lattice. Sf9 tau displayed the lowest microtubule affinity, but conversely the highest cooperative interactions during the envelope formation. Right panels showing the role of tau phosphorylation in CTE-like filament co-assembly. Sf9 tau displayed the highest propensity for CTE-like filament co-assembly, indicating the phosphorylation pattern, but not the total level of phosphorylation, dramatically alters the propensity of full-length tau molecules to enter into the pathological conformation.

**Table 1:** Site-specific phosphorylation of eukaryotic expressed 2N4R tau.

| site | Degree of phosphorylation |  | site | Degree of phosphorylation |  |
| --- | --- | --- | --- | --- | --- |
|  | HEK tau | SF9 tau |  | HEK tau | SF9 tau |
| T17 | 0 | (0.06 ± 0.08) % | T231 | (97.47 ± 0.02) % | (99.81 ± 0.28) % |
| Y18 | 0 | (0.01 ± 0.01) % | S235 | (34.46 ± 37.78) % | (29.58 ± 40.68) % |
| Y29 | 0 | (1.10 ± 1.55) % | S237 | (25.96 ± 0.59) % | (49.17 ± 36.31) % |
| T30 | (0.19 ± 0.27) % | (0.41 ± 0.58) % | S238 | 0 | (0.03 ± 0.05) % |
| T39 | 0 | (0.20 ± 0.28) % | S241 | (17.30 ± 24.47) % | (4.80 ± 6.79) % |
| S46 | (10.72 ± 15.16) % | (1.48 ± 1.93) % | T245 | (0.02 ± 0.04) % | (0.01 ± 0.01) % |
| T50 | (3.19 ± 4.51) % | 0 | S258 | (16.84 ± 23.81) % | 0 |
| T52 | 0 | (0.01 ± 0.02) % | S262 | (0.50 ± 0.70) % | 0 |
| S56 | 0 | (0.01 ± 0.02) % | T263 | (0.16 ± 0.22) % | (0.07 ± 0.09) % |
| S61 | 0 | (0.21 ± 0.30) % | S285 | (0.06 ± 0.08) % | (0.003 ± 0.005) % |
| S68 | (1.27 ± 1.64) % | 0 | S289 | (0.01 ± 0.01) % | (0.01 ± 0.02) % |
| T69 | (0.07 ± 0.10) % | 0 | S293 | (0.72 ± 1.03) % | (2.31 ± 3.27) % |
| T71 | (0.29 ± 0.21) % | 0 | S305 | (0.84 ± 1.04) % | (0.56 ± 0.78) % |
| T76 | 0 | (0.25 ± 0.35) % | Y310 | (0.16 ± 0.04) % | (0.39 ± 0.52) % |
| T102 | (5.70 ± 8.05) % | 0 | S316 | (0.30 ± 0.43) % | 0 |
| T111 | 0 | (0.01 ± 0.01) % | T319 | (24.62 ± 34.81) % | 0 |
| S113 | (0.14 ± 0.19) % | (0.02 ± 0.03) % | S320 | (5.65 ± 8.00) % | 0 |
| T123 | 0 | (0.02 ± 0.03) % | S324 | (1.31 ± 0.17) % | (0.54 ± 0.77) % |
| T169 | (4.55 ± 6.43) % | (1.08 ± 1.53) % | S341 | (0.06 ± 0.09) % | 0 |
| T175 | (3.58 ± 4.57) % | (7.78 ± 4.23) % | S352 | (4.53 ± 6.41) % | (44.70 ± 63.22) % |
| T181 | (72.42 ± 11.34) % | (66.42 ± 8.49) % | S356 | (0.51 ± 0.58) % | (0.76 ± 0.76) % |
| S184 | 0 | (0.37 ± 0.53) % | T361 | (0.03 ± 0.04) % | (0.02 ± 0.02) % |
| S185 | 0 | (0.18 ± 0.25) % | T373 | (29.56 ± 41.80) % | 0 |
| S191 | (0.60 ± 0.85) % | (10.80 ± 15.28) % | T377 | (8.13 ± 11.50) % | 0 |
| S195 | (0.97 ± 1.38) % | (2.54 ± 3.59) % | T386 | 0 | (0.08 ± 0.11) % |
| Y197 | (0.35 ± 0.50) % | 0 | Y394 | (0.63 ± 0.89) % | (3.99 ± 5.64) % |
| S198 | (0.25 ± 0.36) % | (22.99 ± 4.81) % | S396 | (11.53 ± 15.69) % | (29.84 ± 32.75) % |
| S199 | (3.45 ± 4.88) % | (4.82 ± 6.82) % | S400 | (1.54 ± 0.13) % | (7.40 ± 9.85) % |
| S202 | (16.81 ± 15.44) % | (47.06 ± 14.81) % | T403 | (18.32 ± 25.69) % | (10.81 ± 15.29) % |
| T205 | (1.42 ± 2.01) % | (0.99 ± 1.41) % | S404 | (22.30 ± 25.04) % | (60.41 ± 17.79) % |
| S208 | (0.07 ± 0.10) % | (13.76 ± 19.46) % | S409 | 0 | (0.12 ± 0.17) % |
| S210 | (15.87 ± 22.14) % | (42.88 ± 56.19) % | T414 | 0 | (0.12 ± 0.17) % |
| T212 | (0.33 ± 0.47) % | (2.58 ± 2.95) % | S416 | 0 | (1.44 ± 1.69) % |
| S214 | 0 | (0.85 ± 0.87) % | S422 | 0 | (31.71 ± 42.91) % |
| T217 | (7.15 ± 10.08) % | (5.29 ± 1.17) % | T427 | (10.61 ± 14.60) % | (0.83 ± 1.17) % |
| T220 | (0.01 ± 0.01) % | (0.08 ± 0.12) % | S435 | 0 | (0.02 ± 0.03) % |

Average phosphorylation degree:
HEK tau: ~ 9.30 % SF9 tau: ~ 10.58 %

## Acknowledgements

We thank all the members of the MOM lab for their continual input and feedback on this project. The authors thank previous lab members Aileen Lam and Mariah Dacy for generating tau constructs for insect and mammalian expression. The authors thank Armann Andaya (UC Davis Campus Mass Spectrometry Facilities) for the help of mass spectrometry. The authors thank Fei Guo and Bradley Shibata (UC Davis Biological Electron Microscopy Facility) for the help of negative-stain transmission electron microscopy. The work was supported by UC Davis Alzheimer’s Disease Research Center (ADRC) Development Project (P30AG072972) and an Alzheimer’s Association Research Grant: 23AARGD-1022898 (to KMO).

## Author contributions

X.F. and R.J.M conceived the project and designed the research. R.J.M. and K. M. O.-M. secured research funding. X.F performed the research and analyzed the data. K.O. performed the microtubule binding assay for the hill curve and optimized the mammalian tau construct by adding a WPRE element. H.L expressed and purified StrepII-GFP-tau in bacterial and Sf9 cells. X.F and R.J.M wrote the paper. All authors edited the paper.

## Material and Methods

### Plasmids

A Full-length human tau cDNA containing plasmid was purchased from Addgene (16316). Coding sequences of Full-length human 2N4R tau or its truncated fragment dGAE (aa297-391) were amplified by PCR and subsequently cloned into a pET28a vector (Invitrogen) or a pFastBac vector (Invitrogen) or a pAceMam2 vector (GenevaBiotech) containing either an N-terminal StrepII-sfGFP or StrepII-SNAPf by Gibson^13,93^. To increase the gene expression, the pAceMam2 vector was further optimized by adding a woodchuck hepatitis virus post-transcriptional regulatory element (WPRE) downstream of the ORF^94^. All constructs were verified by Sanger sequencing.

### Protein expression and purification

dGAE, StrepII-sfGFP-tau and StrepII-SNAPf-tau in pET28 were expressed in BL21-CodonPlus (DE3)-RIPL cells (Agilent). Briefly, a single colony was inoculated into 50 ml LB Broth medium (Fisher Scientific) and grew overnight at 37 °C, the culture was then diluted 1 to 500 into 500 mL LB Broth medium (Fisher Scientific) and cultured at 37 °C until the optical density at 600 nm (OD600) reached 0.5. Expressions were induced 18 h by 0.4 mM isopropyl-β-d-thiogalactoside (Fisher Scientific) at 24 °C for dGAE and StrepII-sfGFP-tau, for StrepII-SNAPf-tau, the expression was induced 4h by 0.4 mM isopropyl-β-d-thiogalactoside (Fisher Scientific) at 37 °C. Cells were subsequently harvested by centrifuge at 1000×g for 10 min and flash frozen by liquid nitrogen. All frozen cells were stored at −80 °C before purification.

StrepII-sfGFP-tau and StrepII-SNAPf-tau in pFastBac were expressed in insect cells. Briefly, insect Sf9 cells were grown or maintained in Sf-900 II serum-free medium (Thermo Fisher Scientific) at 27 °C. Plasmids DNA of corresponding constructs were transformed into DH10EmBacY competent cells (Geneva Biotech) to generate bacmids. To get baculovirus, 1 × 10^6^ Sf9 cells were transferred to each well of a six-wells plate and subsequently transfected with 1 μg of bacmid DNA by 6 µl of Cellfectin (Thermo Fisher Scientific). Baculovirus-containing cell supernatants (P1) were harvested 7 days after transfection when the entire cell population has become infected. To amplify the baculovirus (P2), 50 ml of Sf9 cells were infected with 50 µl of P1 virus, and the P2 virus was harvested 5 days after infection. To prepare recombinant proteins, 200 ml of Sf9 cells at high density (∼ 2 × 10^6^ cells/ml) were infected by P2 virus (virus: cells = 1:100, v/v) and cultured around 65 h at 27 °C. Cells were subsequently harvested by centrifuge at 1000 × g for 10 min and flash frozen by liquid nitrogen.

StrepII-sfGFP-tau and StrepII-SNAPf-tau in pAceMam2 were expressed in mammalian cells. Briefly, suspension-adapted human embryonic kidney (HEK) cell line FreeStyle 293-F cells (Invitrogen) were grown or maintained in FreeStyle™ 293 Expression Medium (Thermo Fisher Scientific) at 37 °C with a humidified atmosphere of 8% CO_2_. To prepare recombinant proteins, 400 ml of 293-F cells at high density (∼ 1 × 10^6^ cells/ml) were transiently transfected by corresponding plasmid DNA (v_cells_: m_DNA_= 1:1 ml/μg), the transient transfections were mediated by polyethyleneimine (PEI, m_DNA_: m_PEI_= 1:4). The transfected cells were cultured around 72 h at 37 °C with a humidified atmosphere of 8% CO_2_. Cells were subsequently harvested by centrifuge at 1000 × g for 10 min and flash frozen by liquid nitrogen.

To purify the dGAE protein, the frozen pellet 1L culture was thawed on ice and resuspended in 40 ml of wash buffer (WB: 50 mM MES pH 6.0, 10 mM EDTA, 10 mM DTT), freshly supplemented with 1 mM dithiothreitol (DTT), 1 mM phenylmethylsulfonyl fluoride (PMSF), protease inhibitor mix (Promega) and 5 µl of Benzonase Nuclease (Millipore). Cells were lysed by passing through an Emulsiflex C-3 (Avestin) and were centrifuged at 15, 000×g for 20 min at 4 °C to obtain soluble lysate. clarified lysate was filtered through 0.45 μm filters and loaded onto a HiTrap SP HP 5 ml column (GE Healthcare) for cation exchange. Bound proteins were eluted with 30 mL of a linear gradient of 0–1 M NaCl in WB buffer. Protein-containing fractions were pooled and precipitated using 30% (m/v) ammonium sulphate and left on a rocker for 30 min at 4°C. Precipitated proteins were then centrifuged at 20,000 ×g for 35 min at 4°C and resuspended in 2 ml of 10 mM PB (pH 7.4), freshly supplemented with 10 mM DTT, and loaded onto a 60/10 Superdex 200 size exclusion column. Size exclusion fractions were analyzed by sodium dodecyl sulfate-polyacrylamide gel electrophoresis (SDS-PAGE), protein-containing fractions pooled and concentrated to 6 mg/ml using molecular weight concentrators with a cut-off filter of 3 kDa. The dGAE concentration was measured by NanoDrop One (Thermo Fisher Scientific) based on the UV absorbance at 280 nm using an Abs 0.1% of 0.144 and was further confirmed by comparing with the standard bovine serum albumin (BSA) concentrations on SDS-PAGE gel. Purified protein samples were flash frozen in 100 μl aliquots for future use.

To purify StrepII-sfGFP-tau and StrepII-SNAPf-tau, frozen pellets were thawed on ice and resuspended in 40 ml of purification buffer (50 mM Tris–HCl pH 8.0, 150 mMKCH_3_COO, 2 mM MgSO_4_, 1 mM EGTA, 5% glycerol) freshly supplemented with 1mM dithiothreitol (DTT), 1 mM phenylmethylsulfonyl fluoride (PMSF), and protease inhibitor mix (Promega). Resuspended cells were lysed by passing through an Emulsiflex C-3 (Avestin) (for *E. coli* expression), or were homogenized in a dounce homogenizer and lysed by the addition of 1% Triton X-100 and 5 µl of Benzonase Nuclease (Millipore) for 10 min on ice (for insect cell and mammalian cell expression). To obtain soluble lysate, lysed cells were centrifuged at 15, 000×g for 20 min at 4 °C. The clarified lysate was incubated with 10 ml of Strep-Tactin XT resin (IBA) for 1 h at 4 °C and washed with purification buffer, protein was subsequently eluted by elution buffer (purification buffer plus 100 mM biotin, pH 8.0). Eluted proteins were subsequently diluted 20x by IEX Buffer pH 8.0 (Ionic Exchange Buffer: 50 mM Tris-HCl pH 8.0, 2 mM MgCl_2_, 1 mM EGTA and 10% glycerol) and were loaded onto a 5 ml HiTrap Q HP column (Cytiva), bound proteins were eluted with a linear gradient of 0–1 M NaCl in IEX Buffer. Eluted protein was further purified by Superdex 200 columns (Cytiva) in PB buffer. Protein-containing fractions were pooled and supplemented with 10% Glycerol. Protein concentrations were measured by NanoDrop One (Thermo Fisher Scientific) based on the absorbance of the attached fluorophores (StrepII-sfGFP-tau) or the UV absorbance at 280 nm using an Abs 0.1% of 0.550. All protein aliquots were subsequently flash frozen by liquid nitrogen and stored at −80 °C.

### Labeling differentially phosphorylated tau proteins with SNAP tag substrate

A part of differentially phosphorylated StrepII-SNAP-tau proteins were fluorescently labeled with SNAP-Surface 649 (NEB) according to the manufacturer’s protocols. Briefly, differentially phosphorylated StrepII-SNAP-tau proteins were diluted to ∼ 25 μM by purification buffer (50 mM Tris–HCl pH 8.0, 150mM KCH_3_COO, 2 mM MgSO_4_, 1 mM EGTA, 5% glycerol), then incubated with 50 μM of SNAP-Surface 649 on ice for 2 hours. Excess SNAP-Surface 649 was removed by 5 ml HiTrap Q HP column (Cytiva) equilibrated with PB buffer.

### Mass Spectrometry

Protein samples were treated with standard tryptic digestion. Briefly, samples were reduced in 5.5 mM DTT at 56°C for 45 minutes, followed by alkylation with iodoacetamide added to a final concentration of 10 mM for one hour. Subsequently, samples were digested overnight by trypsin at a final mass ratio of 1:50 (enzyme: substrate) at 37°C. The reaction was quenched by flash freezing in liquid nitrogen and the digest was lyophilized. Digest was reconstituted in 0.1% TFA with 10% acetonitrile prior to injection.

Samples were analyzed using a nanoAcquity UPLC system (Waters) connected to the Xevo G2 QTof mass spectrometer. Samples were loaded onto a C18 Waters Trizaic nanotile of 85 μm × 100 mm; 1.7 μm (Waters). Mass spectrometry data were recorded for 60 minutes for each run and controlled by MassLynx 4.2 SCN990 (Waters). Acquisition mode was set to positive polarity under resolution mode. RAW files were processed using Protein Lynx Global Server (PLGS) version 3.0.3 (Waters). For mass spec analysis, data was searched using Protein Lynx Global Server 3.0.3 Expression 2.0 (Waters).

Site-specific phosphorylation degrees were estimated by directly comparing the total ion intensities of a phosphopeptide/peptide pair at certain sites as previous description^54,55^.

### Mass photometry

The mass photometry assays were carried out as previously described^95^. Briefly, to prepare chambers, microscope cover glass (#1.5 24 × 50 mm, Deckgläser) was cleaned by 1 h sonication in Milli-Q H_2_O, followed by another hour sonication in isopropanol, cover glasses were then washed by Milli-Q H_2_O and dried by flame. CultureWell silicone gaskets (Grace Bio-Labs) were cut and washed by Milli-Q H_2_O, CultureWell silicone gaskets were then dried by filtered air and placed onto the freshly cleaned cover glasses providing four independent sample chambers. Samples and standard proteins were diluted in SRP90 buffer (90 mM HEPES, 10 mM PIPES, 50 mM KCH_3_COO, 2 mM Mg(CH_3_COO)_2_, 1 mM EGTA, 10% glycerol, pH = 7.4). The SRP90 buffer was freshly filtered by a 0.22 µm filter before measurement. For calibration, standard proteins BSA (Sigma), Alcohol dehydrogenase (Sigma) and beta-amylase (Sigma) were diluted to 5–50 nM. The non-specific binding events of single molecules on cover glasses were scattered and the masses of these molecules were measured by the OneMP instrument (Refeyn) at room temperature. Data were collected at an acquisition rate of 1 kHz for 100 s by AcquireMP (Refeyn) and subsequently analyzed by DiscoverMP (Refeyn). For each concentration of recombinant proteins, the measurement was performed three times and repeated with two different protein preparations. Results from one representative measurement are shown in the figure.

### Microtubule preparation

Porcine brain tubulin was isolated using the high-molarity PIPES procedure and then labeled with biotin NHS ester or Dylight-405 NHS ester as described previously (https://mitchison.hms.harvard.edu/files/mitchisonlab/files/labeling_tubulin_and_quantifying_labeling_stoichiometry.pdf). Pig brains were obtained from a local abattoir and used within ∼ 4h after death. To polymerize microtubules, 50 µΜ of unlabeled tubulin, 10 µM of biotin-labeled tubulin, and 3.5 µM of Dylight-405-labeled tubulin were incubated with 2 mM of GTP for 20 min at 37 °C. Polymerized microtubules were stabilized by the addition of 20 µΜ taxol and incubated additional 20 min. Microtubules were pelleted at 20,000×*g* by centrifugation over a 150 µl of 25% sucrose cushion and the pellet was resuspended in 50 μl BRB80 (80 mM Pipes (pH 6.8), 1 mM MgCl_2_ and 1 mM EGTA) containing 10 µΜ taxol. The same method was used for microtubules polymerized from recombinant tubulin.

### Total internal reflection fluorescence (TIRF) assays

TIRF chambers were assembled from acid-washed glass coverslips as previously described (http://labs.bio.unc.edu/Salmon/protocolscoverslippreps.html), pre-cleaned slide, and double-sided sticky tape. Chambers were first incubated with 0.5 mg/ml PLL-PEG-biotin (Surface Solutions) for 5–10 min, followed by 0.5 mg/ml streptavidin for 5 min. Microtubules were diluted by BRB80 containing 10 µΜ taxol. Diluted microtubules were flowed into streptavidin-adsorbed flow chambers and incubated for 5–10 min at room temperature for adhesion. To remove unbound microtubules, chambers were subsequently washed twice with SRP90 assay buffer (described above). Purified tau protein was diluted to indicated concentrations in the assay buffer with an oxygen scavenging system composed of PCA/PCD/Trolox. Then, the solution was flown into the glass chamber. To quantify the dwell times, events longer than three pixels were selected and quantified and two independent experiments were performed for each condition.

For the SiR-tubulin assay, chambers were preincubated 10 min at room temperature with 2 µM of SiR-tubulin in SRP90 assay buffer after washing. Then differentially phosphorylated tau proteins were diluted to indicated concentrations with 2 µM SiR-tubulin containing SRP90 assay buffer and were flown into the glass chamber respectively. Movies and images were acquired using a NIS-Elements software-controlled (AR, Version 5.20.2) Nikon Eclipse Ti2 Microscope (1.49 numerical aperture, ×100 objective) equipped with a TIRF illuminator and Andor iXon charge-coupled device electron multiplying camera. Data were analyzed manually using ImageJ (Fiji), and statistical tests were performed in GraphPad Prism 10.

### CTE-like filaments co-assembly assay

CTE-like filaments were assembled by dGAE alone (400 µM) or co-assembled by 99% dGAE (396 µM) plus 1% full-length tau (4 µM), the method was slightly modified from previous publication^50,51,96^. Briefly, 300µl of purified dGAE (400 µM) or 300µl of dGAE (396 µM) plus differentially phosphorylated full-length SNAP-tau proteins (4 µM) were incubated in 10mM PB (phosphate buffer: pH7.4) respectively, freshly supplemented with 10 mM dithiothreitol (DTT), 200mM NaCl, 1 mM phenylmethylsulfonyl fluoride (PMSF), protease inhibitor mix (Promega) and 0.01 % NaN_3_. Then the mixed samples were agitated by a MS 3 digital shaker (IKA) at 37 °C (750 rpm) for 96 hours. 40µl of samples were taken at corresponding timepoints and were subsequently flash frozen by liquid nitrogen and stored at −80 °C for further using.

### Analysis co-assembly efficiency by SDS-PAGE

40µl of co-assembled samples froze at different timepoints were thawed and ultracentrifuged 100,000 rpm at room temperature for 20 min by using an Optima MAX-XP ultracentrifuge with a TLA-100.1 rotor (Beckman Coulter). Supernatants were transferred to a 500µl Eppendorf tubes, pellet were resuspended with 40µl of PB buffer (pH7.4). Then the 12µl of supernatants and pellets were loaded and run on the sodium dodecyl sulfate polyacrylamide gel electrophoresis (SDS-PAGE), then the gels were stained with Coomassie blue. The protein intensities of full-length tau proteins in both supernatants and pellets at corresponding timepoints were subsequently measured and compared by Image J.

### Negative-stain TEM

Samples from pellets after 96 hours agitation were centrifuged 15, 000 rpm at room temperature for 15min, the supernatants were discarded, then the pellets were washed 3 times with PB buffer (pH7.4) and were resuspended with 40µl of PB buffer. 5μl of samples from resuspended pellets were incubated on a discharged carbon-coated grid at room temperature for 1 min, excess solution was subsequently removed by filter paper, the grids were negatively stained 5 times with 5μl of 0.75% Uranyl Formate, 30s per time. The grids were left to air dry for 10 min and then were examined on a JEOL 1230 Transmission Electron Microscope.

